# Oncogenic mutation or overexpression of oncogenic KRAS or BRAF is not sufficient to confer oncogene addiction

**DOI:** 10.1101/2020.05.26.117911

**Authors:** Reina E. Ito, Chitose Oneyama, Kazuhiro Aoki

## Abstract

Oncogene addiction is a cellular property by which cancer cells become highly dependent on the expression of oncogenes for their survival. Oncogene addiction can be exploited to design molecularly targeted drugs that kill only cancer cells by inhibiting the specific oncogenes. Genes and cell lines exhibiting oncogene addiction, as well as the mechanisms by which cell death is induced when addicted oncogenes are suppressed, have been extensively studied. However, it is still not fully understood how oncogene addiction is acquired in cancer cells. Here, we take a synthetic biology approach to investigate whether oncogenic mutation or oncogene expression suffices to confer the property of oncogene addiction to cancer cells. We employed human mammary epithelium-derived MCF-10A cells expressing the oncogenic KRAS or BRAF. MCF-10A cells harboring an oncogenic mutation in a single-allele of KRAS or BRAF showed weak tumorigenic activity, but no characteristics of oncogene addiction. MCF-10A cells overexpressing oncogenic KRAS demonstrated the tumorigenic activity, but MCF-10A cells overexpressing oncogenic BRAF did not. Neither cell line exhibited any oncogene addiction properties. These results indicate that the introduction of oncogenic mutation or the overexpression of oncogenes is not sufficient for cancer cells to acquire oncogene addiction, and that oncogene addiction is not associated with tumorigenic potential.

## Introduction

Most human cancers develop over a long period of time, from a few years to several decades, as mutations accumulate in various proto-oncogenes and tumor suppressor genes ^1,2^. During this process, cancer cells rewire the intracellular signal transduction system by accumulating mutations and epigenetic changes, and consequently acquire the characteristics of malignant tumors. On the other hand, it is well-established that the overexpression of oncogenes suffices for the neoplastic transformation of non-cancerous cells *in vitro* and *in vivo*, resulting in infinite proliferation, anchorage independence, and angiogenesis ^3–5^. Therefore, properties that can be acquired over a long period of time appear to be different from the tumorigenesis induced by the proto-oncogenes/tumor suppressor genes activation/inactivation.

Oncogene addiction (or oncogene pathway addiction) is a characteristic of cancer cells in which malignant cells are dependent for their proliferation and survival on a particular proto-oncogene and/or tumor suppressor gene ^6,7^. Thus, the proliferation and survival of oncogene-addicted cancer cells are dramatically impaired by suppression of the oncogenes. For example, the inhibition of addicted oncogenes with RNAi or small chemical inhibitors causes apoptosis in oncogene-addicted cancer cells, but not in other cells (Fig. 1), thereby providing a rationale for molecularly targeted therapy ^8^. Imatinib (Gleevec), a BCR-ABL1 kinase inhibitor, and Gefitinib (Iressa), an EGFR inhibitor, are typical examples of drugs successfully targeted to the appropriate molecules and are effective for the treatment of chronic myeloid leukemia (CML) and non-small cell lung cancer, respectively ^9^. Several molecular mechanisms by which cancer cells die through acute inhibition of addicting oncogenes selectively required for survival have been reported, including oncogene shock, oncogene amnesia, genetic streaming, synthetic lethality, and others ^10,11^. However, little is know about how and when the property of oncogene addiction is acquired, and which oncogene(s) is prone to cause oncogene addiction, although the phenomenon has been reported to involve epigenetic DNA changes that accompany the development of cancer ^12^.

**Figure 1.**
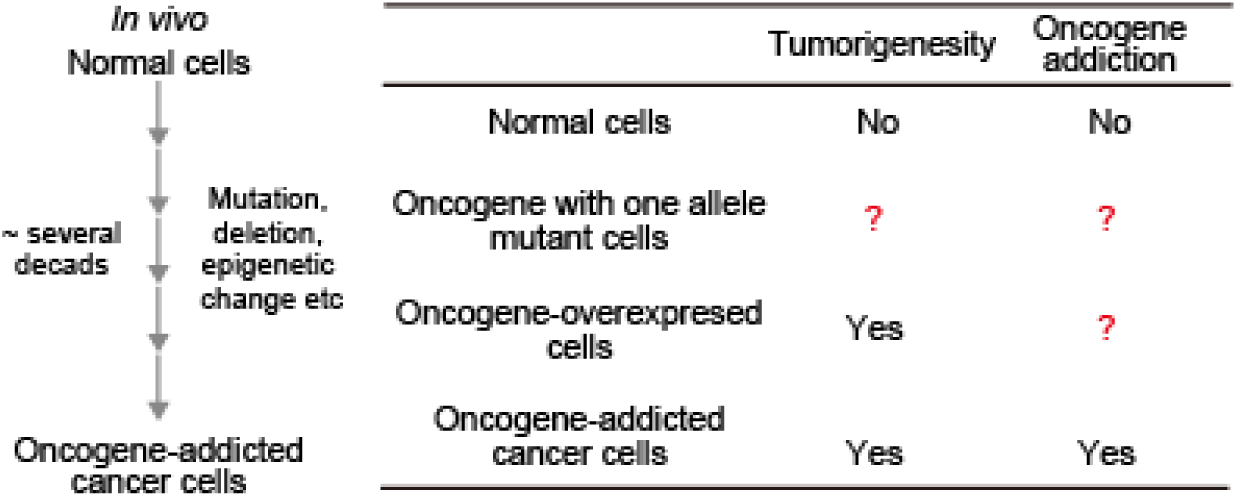
Tumorigenicity and oncogene addiction. (A) *In vivo* process by which cancer cells acquire oncogene addiction. (B) The table compares different type of cells with respect to tumorigenicity and oncogene addiction.

The Ras-ERK signaling pathway plays a pivotal role in a wide range of cell functions such as cell proliferation, differentiation, and survival, but also plays a key role in tumorigenesis ^13,14^. Indeed, the *KRAS* gene is the second-most frequently mutated gene in human cancers, after the *p53* gene, and the *BRAF* gene is also frequently mutated in melanoma and colorectal cancer ^2^. KRAS- or BRAF-mutated cancer cells also exhibit oncogene addiction. Suppression of the expression of mutated KRAS by antisense or siRNA caused cell cycle arrest and apoptosis in KRAS-mutated cultured cancer cell lines, and epithelial-mesenchymal transition (EMT) was closely associated with KRAS dependency ^15,16^. Knockdown by RNAi or treatment with a BRAF-selective inhibitor leads to the inhibition of cell proliferation and survival in BRAF-mutated cancer cell lines ^17–20^. However, these results were obtained by using cell lines established from human patients, it is impossible to trace when and how oncogene addiction is acquired. Interestingly, the expression of oncogenic HRAS or KRAS has been shown to induce tumor formation in a doxycycline-inducible *in vivo* tumorigenesis model, and withdrawal of the drug resulted in tumor shrinkage ^21,22^. However, Chin et al. also showed that these cells do not alter their growth rate regardless of doxycycline treatment *in vitro* ^*21*^. Thus, it remains unclear whether oncogene addiction is achieved by the acquisition of tumorigenic properties through expression of the *KRAS* or *BRAF* oncogenes. In this study, we examined whether an oncogenic mutation in a single allele of *KRAS* or *BRAF* or overexpression of *KRAS* or *BRAF* oncogenes was sufficient to induce oncogene addiction.

## Results

### *In vitro* characterization of MCF-10A cells harboring a single allele mutation of KRAS G12V or BRAF V600E

The human normal mammary gland-derived MCF-10A cell lines were used in this study. The MCF-10A cells were spontaneously immortalized without defined factors ^23^. They are not tumorigenic, i.e., they are not able to grow under anchorage-independent conditions or to form tumors when injected subcutaneously into nude mice ^24^.

To reconstitute oncogene addiction, we first obtained MCF-10A cells harboring KRAS G12V or BRAF V600E mutation, which were generated by genome editing with adeno-associated virus (hereafter referred to as KRAS G12V/+ or BRAF V600E/+ cells) ^24,25^. Mutation of a single allele of KRAS G12V and BRAF V600E was confirmed by direct sequencing (Fig. S1A). KRAS G12V/+ cells grew as efficiently as parental MCF-10A cells onto two-dimensional dishes, showing islands of densely packed cells (Fig. 2A). Meanwhile, BRAF V600E/+ cells exhibited more scattered and fibroblastic morphology (Fig. 2A). We next evaluated anchorage-independent colony formation in soft agar, which is a feature of transformation ^26^. Seven days after seeding in the soft agar, the parental MCF-10A cells hardly proliferated, whereas KRAS G12V/+ cells grew slowly and formed small spheroids (Fig. 2B). Under the same condition, BRAF V600E/+ cells formed larger and more spheroids than KRAS G12V/+ cells (Fig. 2B), suggesting an increase in transformation activity in BRAF V600E/+ cells. Despite the smaller size of colonies, the number of colonies in KRAS G12V/+ cells was comparable to that of BRAF V600E/+ cells 4 weeks after seeding in the soft agar (Fig. S1B and S1C). Consistent with these data, BRAF V600E/+ cells showed significantly higher basal phosphorylation of ERK, downstream of KRAS and BRAF, under normal and serum-starved conditions (Fig. 2C and 2D). Parental and KRAS G12V/+ cells responded well to EGF stimulation, while BRAF V600E/+ cells demonstrated less sensitivity to EGF, probably because of its higher basal activity (Fig. 2C and 2D). The expression levels of KRAS and BRAF showed no substantial changes in these cell lines (Fig. S1D), suggesting that the differences between parental cells and the mutant cell lines were attributable to the increased activity of KRAS and BRAF. These results indicated that a single allele mutation of BRAF V600E in MCF-10A cells enhances *in vitro* tumorigenic activity, while a single allele KRAS G12V results in cells with similar or slightly increased *in vitro* transformation activity.

**Figure 2.**
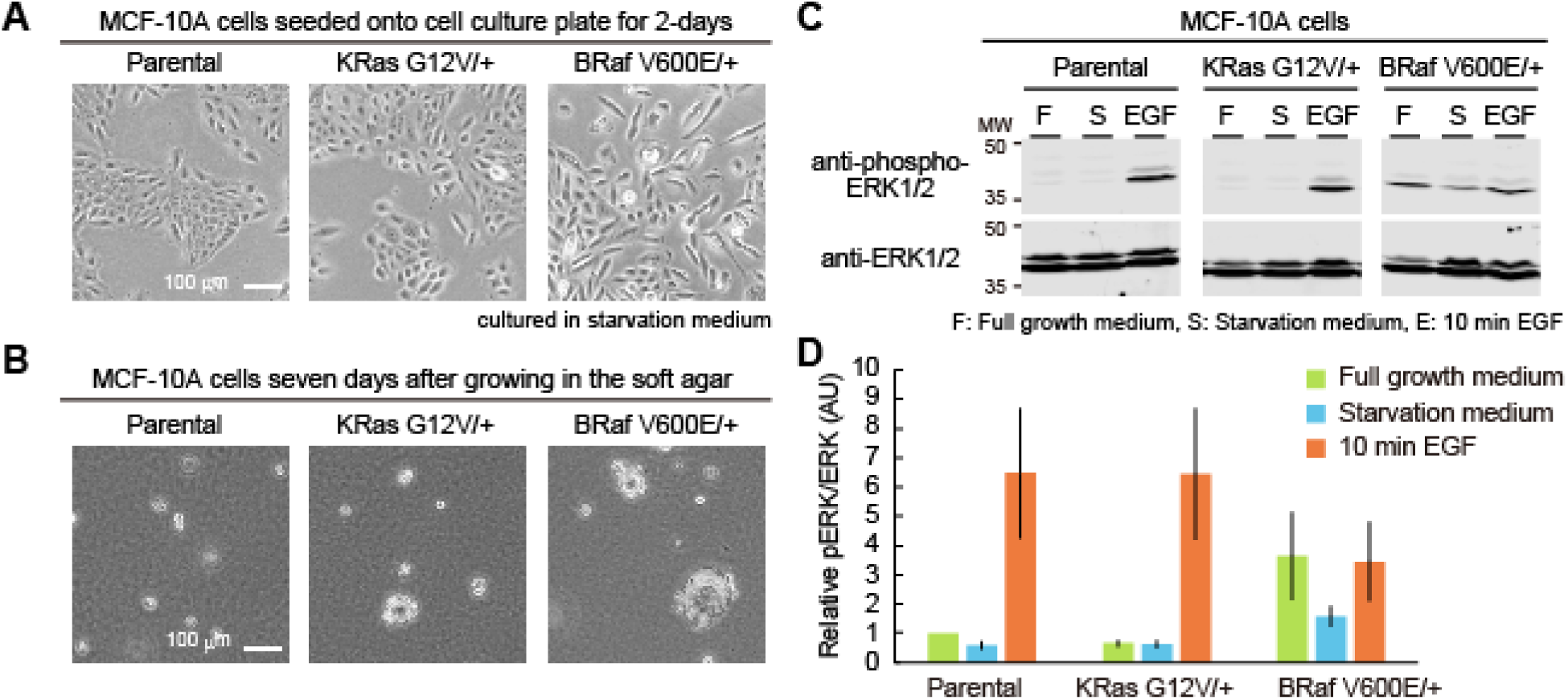
Characterization of MCF-10A cells harboring a single allele mutation of KRAS G12V or BRAF V600E. (A and B) Morphology of the indicated MCF-10A cells seeded onto a cell culture dish for two days (A) or seeded in soft agar for seven days (B). (C) Western blot analysis from the parental MCF-10A, KRAS G12V/+, and BRAF V600E/+ cell lines under the indicated conditions. (D) Quantification of ERK phosphorylation in panel C. Relative pERK/ERK values normalized by parental MCF-10A cells cultured in full growth medium condition are shown with the SD (n > 3).

### Evaluation of oncogenic KRAS or BRAF addiction in MCF-10A cells harboring a single allele mutation of KRAS G12V or BRAF V600E

We next quantified to what extent cells were addicted by the expression of *KRAS* or *BRAF* oncogene. For this purpose, the effect of KRAS or BRAF ablation on cell growth was examined with crystal violet staining and RNA interference (RNAi) ^16,27^. In addition, to reduce the off-target effect of RNAi, siPOOLs were used to deplete KRAS and BRAF; siPOOLs dilute the off target effects by pooling multiple siRNAs against the target genes ^28^.

As a control, we used two lung cancer-derived cell lines, A549 (homozygous KRAS G12S mutation; KRAS non-addicted) and H358 (heterozygous KRAS G12C mutation; KRAS-addicted), and melanoma-derived cell line, A375 (homozygous BRAF V600E mutation; BRAF-addicted) ^16^. The knockdown of KRAS or BRAF with siPOOLs was confirmed in these cell lines (Fig. S2A). A549 cells showed modest inhibition of cell growth by KRAS depletion (Fig. 3A and 3B). H358 and A375 cells exhibited strong suppression of cell growth by KRAS and BRAF ablation with the siPOOLs (siRNA), respectively (Fig. 3A and 3B). ERK phosphorylation seemed not to be correlated with the extent of cell growth inhibition induced by siPOOLs-mediated gene knockdown, in contradiction to the earlier report ^16^. The inhibition of cell growth was observed even under a full growth condition for the cancer cell lines; these cells were indeed addicted to the expression of the KRAS or BRAF oncogene for their cell growth.

**Figure 3.**
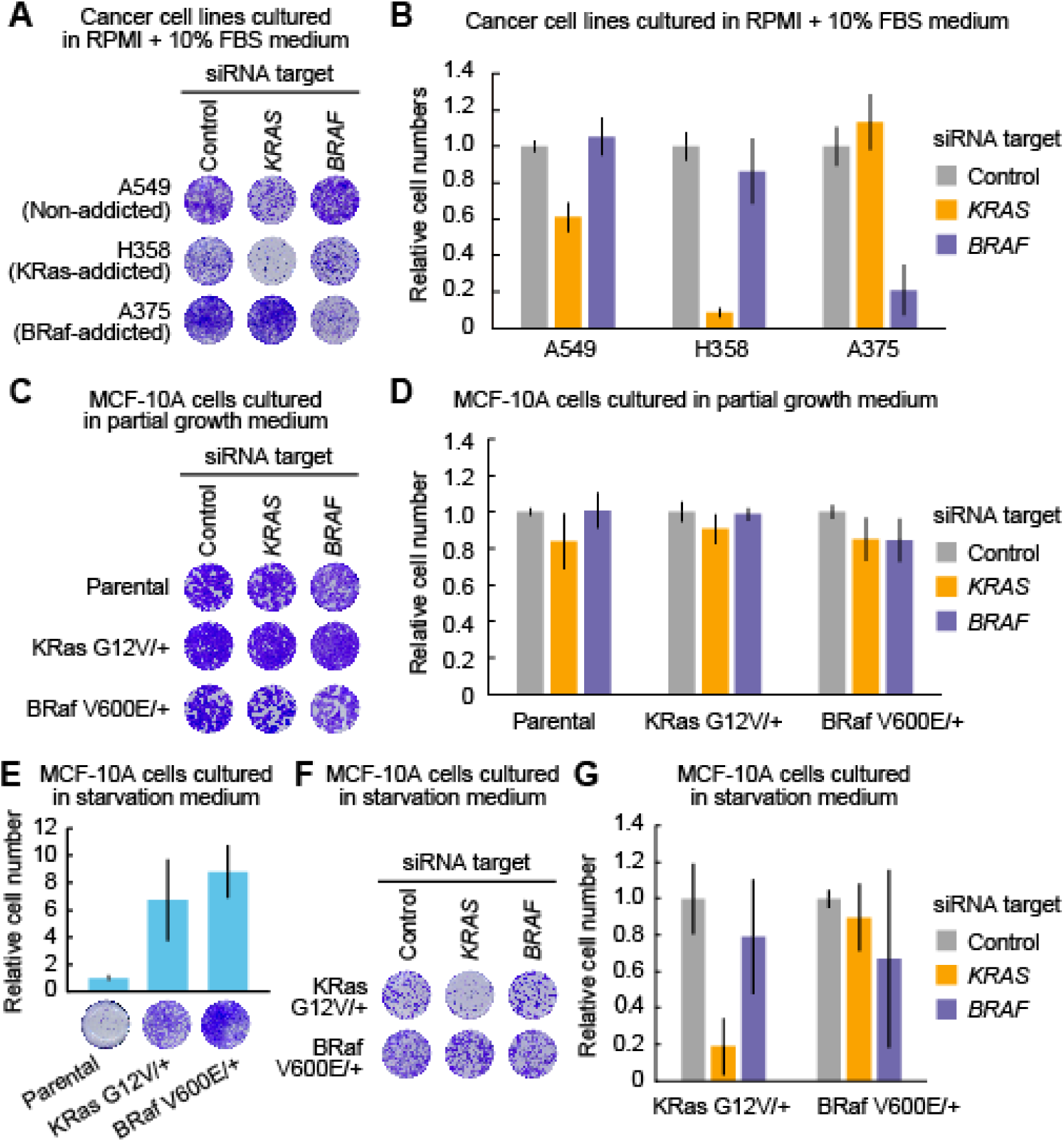
Evaluation of oncogenic KRAS or BRAF addiction in MCF-10A cells harboring a single allele mutation of KRAS G12V or BRAF V600E. (A and B) Cell growth assays following siRNA-mediated negative control, KRAS, or BRAF ablation in the A549, H358, and A375 cell lines. Four days after transfection with siRNA, relative cell densities were determined by crystal violet staining. Representative 96-well plates are shown (A). The mean relative cell number is shown with the SD (n = 2) (B). (C and D) Cell growth assays following siRNA-mediated negative control, KRAS, or BRAF ablation in the parental MCF-10A, KRAS G12V/+, and BRAF V600E/+ cell lines grown in the partial growth medium. Three days after transfection with siRNA, relative cell densities were determined by crystal violet staining. Representative 96-well plates are shown (C). The mean relative cell number is shown with the SD (Parental, n = 4; KRAS G12V/+, n = 4; BRAF V600E/+, n=3) (D). (E) Cell growth assays in the parental MCF-10A, KRAS G12V/+, and BRAF V600E/+ cell lines grown in starvation medium for four days with the indicated siRNA. Relative cell densities were determined by crystal violet staining. Representative 96-well plates are shown (lower). Mean relative cell numbers are shown with the SD (Parental, n = 3; KRAS G12V/+, n = 9; BRAF V600E/+, n=8) (upper). (F and G) Cell growth assays following siRNA-mediated negative control, KRAS, or BRAF ablation in the KRAS G12V/+ and BRAF V600E/+ cell lines grown in starvation medium. Four days after transfection with siRNA, relative cell densities were determined by crystal violet staining. Representative 96-well plates are shown (F). The mean relative cell number is shown with the SD (n = 8) (G).

We then examined whether KRAS G12V/+ and BRAF V600E/+ cells were addicted to their oncogenes. First, siPOOLs were introduced into parental, KRAS G12V/+, and BRAF V600E/+ MCF-10A cells cultured in “full growth medium”. However, we were unable to reproducibly quantify the effect of RNAi on KRAS or BRAF ablation and cell growth. Therefore, we used “partial growth medium”, which contained DMEM and serum without the supplements included in the “full growth medium”. Under the culture condition with the partial growth medium, MCF-10A cells grew and proliferated slowly, making it easier to assess the depletion of KRAS and BRAF (Fig. S3A). Parental MCF-10A cells did not show any detectable change in cell growth by the knock-down of the *KRAS* or *BRAF* gene (Fig. 3C and 3D). We found that neither KRAS nor BRAF depletion resulted in substantial cell growth in KRAS G12V/+ and BRAF V600E/+ cells (Fig. 3C and 3D), indicating that cell growth in these cells was not dependent on the expression of the KRAS or BRAF oncogene. Meanwhile, the oncogenic mutation in a single allele of *KRAS* or *BRAF* leads to sustained proliferative signaling ^2^, enabling MCF-10A cells to grow in the culture medium without EGF and serum, namely, the “starvation medium” (Fig. 3E). Interestingly, the growth factor independence was derived from KRAS expression in KRAS G12V/+ cells, whereas it was not derived from BRAF expression in BRAF V600E/+ cells (Fig. 3F and 3G). In sum, we concluded that KRAS G12V/+ cells and BRAF V600E/+ cells were not addicted to the oncogene, even though these cells acquired the ability to grow without growth factors.

### *In vitro* characterization of MCF-10A cells overexpressing KRAS G12V or BRAF V600E

Next, we examined whether overexpression of KRAS G12V or BRAF V600E induced the property of oncogene addiction. The FLAG-tagged KRAS G12V or BRAF V600E oncogene was introduced into parental MCF-10A cells through lentivirus, producing MCF-10A cells over-expressing KRAS G12V or BRAF V600E (hereinafter referred to KRAS G12V OE or BRAF V600E OE cells)(Fig. S4A and 4B). During the course of experiments, we recognized that long-term culture of BRAF V600E OE cells reduced the expression levels of BRAF V600E and ERK phosphorylation levels (Fig. S4B), probably due to the adaptation through a reduction of BRAF V600E expression and/or negative feedback mechanisms ^29^. Thus, we referred to early-passage (< 1 week from the establishment of cell lines) and late-passage (> 1 week) cells as BRAF V600E OE early cells and BRAF V600E OE late cells, respectively.

The morphology of KRAS G12V OE cells cultured on the plastic dish was scattered and fibroblastic, whereas BRAF V600E OE late cells exhibited a typical epithelial cell shape to the same extent that the empty vector-introduced control cells did (Fig. 4A). KRAS G12V OE cells displayed rapid cell growth in soft agar, forming large colonies one week after seeding (Fig. 4B). Raptured spheroids were observed two weeks after seeding (Fig. S4C), and finally, many small colonies were observed four weeks after seeding (Fig. S4D). BRAF V600E OE late cells showed no anchorage-independent growth in soft agar, but BRAF V600E OE early cells formed small colonies in soft agar after one week (Fig. 4B), large colonies at two weeks that were equal in size to those in anchorage-independent growth of BRAF V600E/+ cells for 4-weeks (Fig. S4C), and finally, a small number of larger colonies at 4 weeks after seeding (Fig. S4D). Consistent with the change in cell morphology and anchorage-independent growth, strong ERK phosphorylation was maintained in KRAS G12V OE cells under all conditions (Fig. 4C and 4D). Despite the higher basal ERK phosphorylation in BRAF V600E OE early cells (Fig. S4B), BRAF V600E OE late cells showed comparable or slightly higher levels of ERK phosphorylation in comparison to control cells (Fig. 4C and 4D).

**Figure 4.**
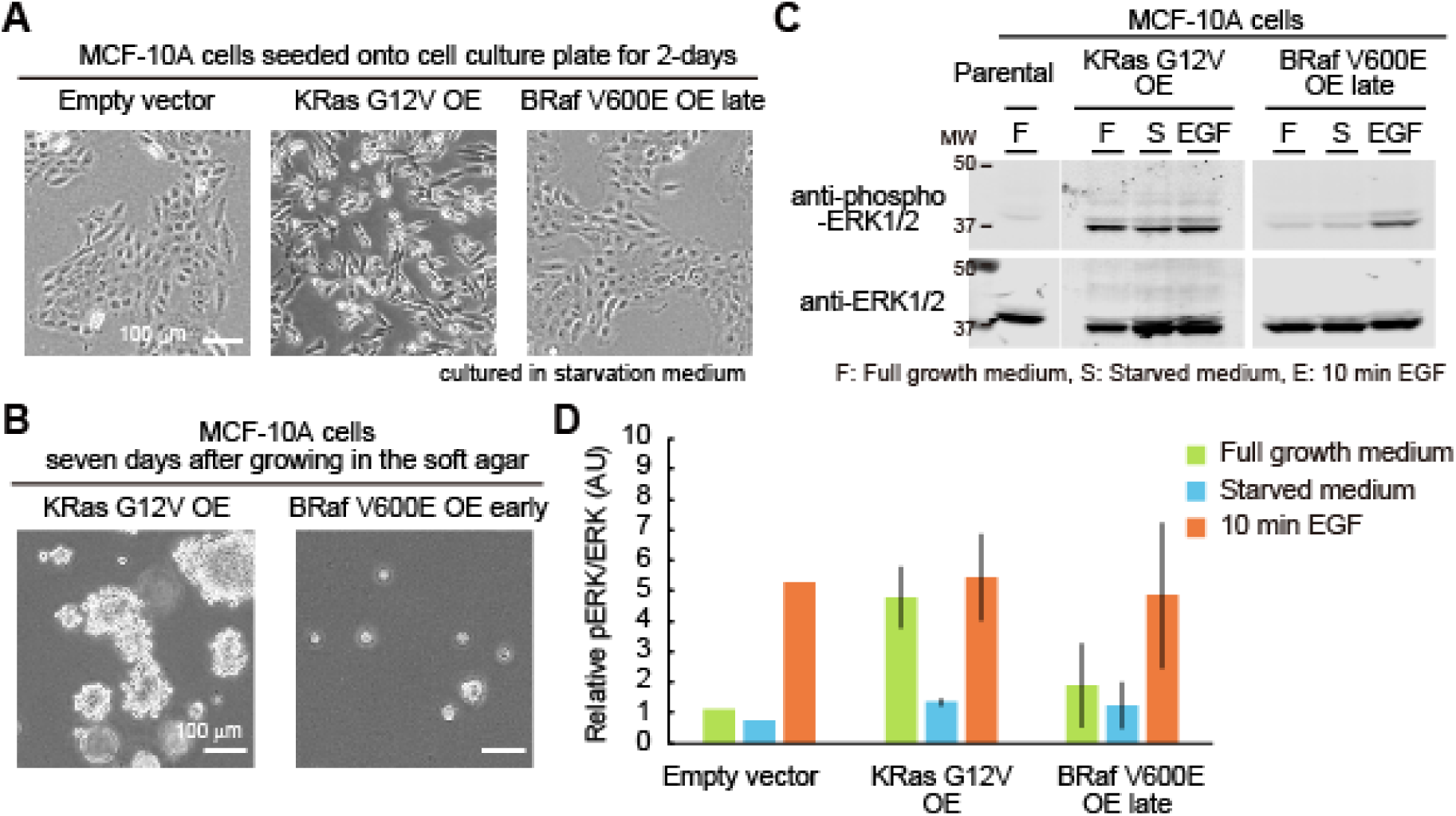
Characterization of MCF-10A cells overexpressing KRAS G12V or BRAF V600E. (A and B) Morphology of the indicated MCF-10A cells seeded onto a cell culture dish for two days (A) or seeded in soft agar for seven days (B). (C) Western blot analysis of the parental MCF-10A, KRAS G12V OE, and BRAF V600E OE late cell lines under the indicated conditions. (D) Quantification of ERK phosphorylation in panel C with empty vector expressing MCF-10A. Relative pERK/ERK values normalized by the parental MCF-10A cells under the full growth medium condition are shown with the SD (Empty vector, n = 1; KRAS G12V OE, n = 5; BRAF V600E OE late, n = 5).

### Evaluation of oncogenic KRAS or BRAF addiction in MCF-10A cells overexpressing KRAS G12V or BRAF V600E

Finally, we quantified the effect of depletion of KRAS and BRAF with siPOOLs (siRNA) on cell growth in KRAS G12V OE and BRAF V600E OE late cells, respectively. The knockdown efficiency of siPOOLs targeting KRAS and BRAF was confirmed by western blotting (Fig. S5A and S5B). To our surprise, KRAS or BRAF ablation did not reduce the cell growth rate in KRAS G12V OE or BRAF V600E OE late cells cultured in the partial growth medium (Fig. 5A and 5B). Like KRAS G12V/+ cells, KRAS G12V OE cells demonstrated growth factor independence (Fig. 5C), and this was dependent on the expression of KRAS (Fig. 5D and 5F). BRAF V600E OE late cells showed a modest increase in cell growth under the serum starvation condition (Fig. 5C). Taken together, the overexpression of KRAS G12V or BRAF V600E enhanced more or less *in vitro* tumorigenicity, but it did not suffice to induce oncogene addiction in MCF-10A cells.

**Figure 5.**
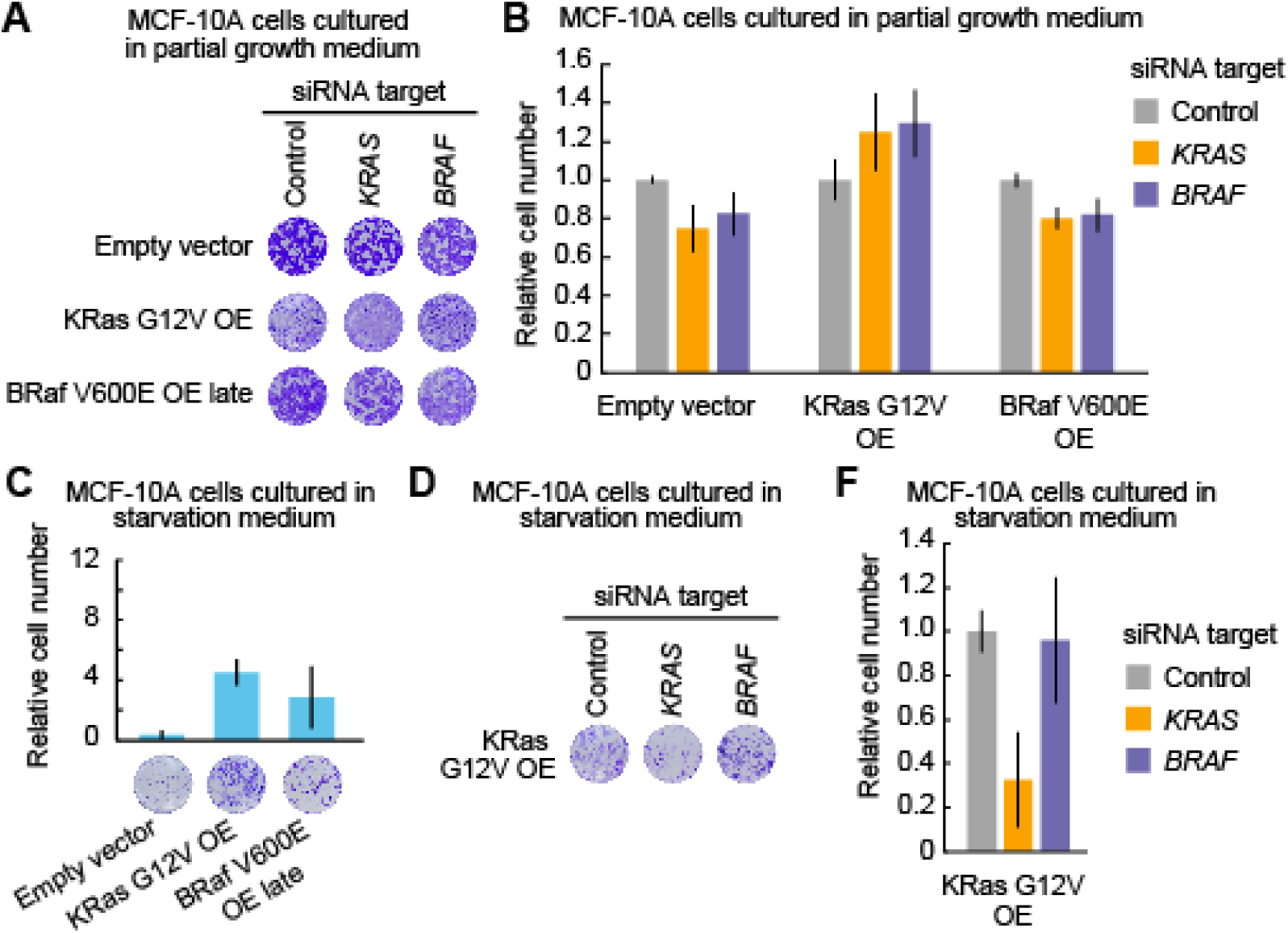
Evaluation of oncogenic KRAS or BRAF addiction in MCF-10A cells overexpressing KRAS G12V or BRAF V600E. (A and B) Cell growth assays following siRNA-mediated negative control, KRAS, or BRAF ablation in parental MCF-10A, KRAS G12V OE, and BRAF V600E OE late cell lines grown in the partial growth medium. Three days after transfection with siRNA, relative cell densities were determined by crystal violet staining. Representative 96-well plates are shown (A). The mean relative cell number is shown with the SD (n = 4) (B). (C) Cell growth assays in empty vector-introduced MCF-10A, KRAS G12V OE, and BRAF V600E OE late cell lines grown in the starvation medium for four days with the indicated siRNA. Relative cell densities were determined by crystal violet staining. Representative 96-well plates are shown (lower). Mean relative cell number are shown with the SD (empty vector control, n = 3; KRAS G12V OE, n =8; BRAFV600E OE, n = 8) (upper). (D and E) Cell growth assays following siRNA-mediated negative control, KRAS, or BRAF ablation in the KRAS G12V OE cell line grown in the starvation medium. Four days after transfection with siRNA, relative cell densities were determined by crystal violet staining. Representative 96-well plates are shown (D). The mean relative cell number is shown with the SD (n = 8) (E).

## Discussion

In this study, we examined the association between oncogene addiction and tumorigenicity (Fig. 6A). In the human normal mammary gland-derived MCF-10A cell lines, an oncogenic mutation in a single allele of the *KRAS* or *BRAF* gene induced modest anchorage-independence, proliferative capacity, and phosphorylation of ERK, while the cells did not exhibit oncogene addiction. Similarly, MCF-10A cells overexpressing the oncogenic KRAS G12V or BRAF V600E protein demonstrated several properties of tumorigenicity, but did not show any signs of oncogene addiction. From these results, we conclude that, at least in the *KRAS* or *BRAF* gene of MCF-10A, the introduction of an oncogenic mutation or overexpression of an oncogene does not ensure the acquisition of oncogene addiction, and the properties of tumorigenicity are not necessarily coupled with oncogene addiction.

**Figure 6.**
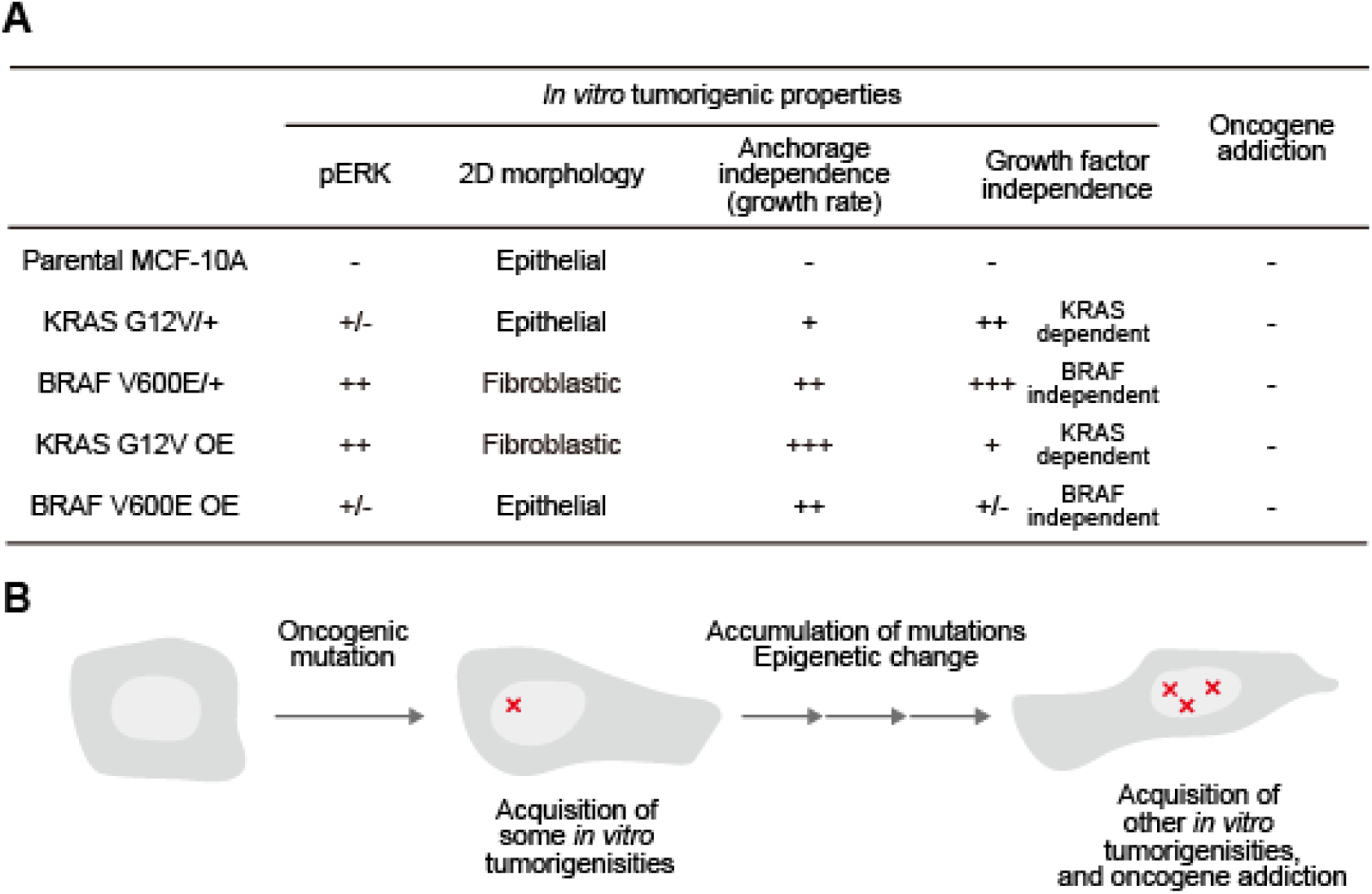
Model for the acquisition of oncogene addiction. (A) A table summarizing the results of this study. (B) Schematic diagram showing how oncogene addiction is acquired.

Why was the oncogenic mutation or over-expression of *KRAS* or *BRAF* in MCF-10A cells not sufficient to induce oncogene addiction? A reasonable possibility is that, after the introduction of the first oncogene mutation *in vivo*, tumor cells accumulate the oncogenes and/or tumor suppressor gene mutations with epigenetic alterations over a long period of time, gradually acquiring oncogene addiction (Fig. 6B). Indeed, sensitivity to BRAF and MEK inhibitors, a feature of BRAF addiction, has been associated with distinct phenotype plasticity of the differentiation state and global alterations in gene expression programs in BRAF-mutated melanomas ^30–32^. Further, it has been reported that a gene expression pattern associated with epithelial-mesenchymal transition (EMT) correlates with KRAS addiction ^16^. Malignant tumor cells exhibiting the property of oncogene addiction may undergo environmental changes *in vivo* that render the cells oncogene-addicted. In other words, the *in vitro* culture of MCF-10A cells may be insufficient to cause genetic and/or epigenetic alteration, leading to oncogene addiction. It would be of critical importance to identify such an environment or stimuli in order to enhance the effects of molecularly targeted drugs. Another possibility is that oncogene addiction may be tissue-specific; thus mammary gland-derived MCF-10A cells may be inherently incapable of acquiring KRAS or BRAF addiction. Oncogenic KRAS or BRAF addiction has been found in lung and colon cancers and melanomas, where *KRAS* or *BRAF* is frequently mutated ^16^. Human breast cancers rarely show *KRAS* and *BRAF* mutations, whereas mutations in genes that activate the PI3K-Akt pathway have been reported ^33^. Interestingly, Her2 amplification and PIK3CA^H1047R^-positive breast cancer exhibited PI3K dependency in a mouse model ^34^, suggesting that genes that are strongly involved in tumorigenesis in a specific tissue are also likely to show oncogene addiction.

In this study, we observed phenotypic differences in MCF-10A cells expressing oncogenic KRAS or BRAF. These differences may have been due to differences in the expression levels and/or differences between KRAS G12V and BRAF V600E. Significant increases in ERK phosphorylation and tumorigenic activity were not observed in KRAS G12V/+ cells (Fig. 2), whereas both changes were observed in KRAS G12V OE cells, which expressed oncogenic KRAS G12V at approximately 20-fold the level of endogenous KRAS (Fig. S4A). Interestingly, neither case demonstrated an oncogene-induced senescence phenotype. It has been reported that cellular senescence is induced when the expression of HRAS G12V is twice that of endogenous RAS in MCF-10A cells and human mammary epithelial cells/hTERT ^35^. Moreover, Caco-2 cells that express low levels of oncogenic KRAS show cellular senescence ^36^. These results suggest that, compared to endogenous KRAS, an approximately two-fold higher expression of oncogenic KRAS is required to induce cellular senescence, and cellular senescence will not be induced by substantially higher or lower levels of expression. With respect to BRAF, our results showed that the expression of BRAF V600E from the endogenous gene locus enhanced ERK phosphorylation and tumorigenicity (Fig. 2), but these characteristic features were gradually diminished when BRAF V600E was overexpressed, and eventually, cells with low BRAF V600E expression were selected (Fig. 4 and Fig. S4). Although BRAF V600E-induced cellular senescence has not been reported in mammary epithelial cells, melanocytes and fibroblasts have been reported to show cellular senescence by overexpression of BRAF V600E ^37^. It is reasonable to assume that overexpression of BRAF V600E induced cellular senescence, thereby leading to the selection of cells with low BRAF V600E expression. However, the susceptibility of oncogene-induced senescence is cell type-dependent. In fibroblasts, overexpression of oncogenic KRAS induces senescence ^38^, while endogenous KRAS G12D expression enhances cell proliferation ^39^. In the future, a more quantitative investigation will be needed to reveal the relationship between the expression level of oncogenes and the consequence of tumorigenicity and cellular senescence.

Future studies should focus on the development of an experimental system for the acquisition of oncogene addiction in various cell lines. It has been reported that overexpression of Myc followed by suppression leads to apoptosis as a model of oncogene addiction ^40^. However, it is technically difficult to trace the process by which cells acquire oncogene addiction in an *in vivo* model. Therefore, it will be necessary to develop an *in vitro* experimental system for the acquisition of oncogene addiction. Understanding what environmental changes lead to oncogene addiction and what state changes the cell undergoes in the process will be important in augmenting the effects of molecularly targeted drugs.

## Materials and methods

### Plasmids, reagents, and antibodies

The plasmids used in this study are listed as follows: pCSIIneo-MCS, pCSIIneo-Flag-BRafV600E, pCSIIneo-Flag-KRasG12V, and pCSIIbsr-Flag-BRafV600E. Plasmids were constructed in accordance with the standard molecular biology methods. The antibodies used for western blot and immunofluorescence analyses are as follows: phospho-anti-Erk1/2 (Thr202/Tyr204) (E10) and anti-Erk1/2 (137F5) were from Cell Signaling Technology; anti-KRAS (clone 3B10-2F2) was from Sigma; anti-Raf-B (F-7) was from Santa Cruz Biotechnology.

### Cell lines

MCF-10A parental cells, MCF-10A BRAF V600E/+ cells, and MCF-10A KRAS G12V/+ cells (catalog numbers HD PAR-003, HD101-012, and HD101-004) were purchased from Horizon Discovery. KRAS G12V OE and BRAF V600E OE were established through lentivirus-mediated gene transfer into the parental MCF-10A cells. In brief, the lentiviral pCSIIneo or pCSIIbsr vectors were transfected into Lenti-X 293T cells (Clontech) together with the packaging plasmid psPAX2 (a gift from Dr. D. Trono, Addgene plasmid #12260), and pCMV-VSV-G-RSV-Rev (a gift of Dr. Miyoshi, RIKEN, Japan) by using the linear polyethyleneimine “Max” MW 40,000 (Polyscience). After two days, MCF-10A parental cells were cultured in the virus-containing media in the presence or absence of 8 μg/mL polybrene for 3-4 hrs. Two days after infection, the cells were selected by at least one-week treatment with 150 ug/ml G418 or 10 μg/mL blasticidin (InvivoGen, San Diego, CA). Bulk populations of selected cells were used in this study. An empty vector, pCSIIneo-MCS, was used as a control. All cell lines were maintained at 37°C under 5% CO2 with antibiotics.

MCF-10A cell lines were maintained in the full growth medium, which consisted of DMEM/F12 (1:1) (Cat#11330-032, Gibco) supplemented with 5% horse serum (Cat#16050-122, Invitrogen), 10 mg/ml insulin (Cat#12878-44, Nacalai Tesque), 0.5 mg/ml hydrocortisone (Cat#1H-0888, Invitrogen), 100 ng/ml cholera toxin (Cat#101B, List Biological Laboratories), 20 ng/ml hEGF (Cat#AF-100-15, PeproTech), and 1% penicillin/streptomycin (Cat#26253-84, Nacalai Tesque). For some experiments, partial growth medium and starvation medium were used; the former contained DMEM/F12 (1:1) supplemented with 5% horse serum and 1% penicillin/streptomycin, and the latter consisted of DMEM/F12 (1:1) supplemented with 2% horse serum, 10 mg/ml insulin, 0.5 mg/ml hydrocortisone, 100 ng/ml cholera toxin, and 1% penicillin/streptomycin.

The A549, H358 (CI-H358), and A375 cell lines were purchased from the American Type Culture Collection. H358 cell lines were maintained in RPMI 1640 media (ATCC modification) (Cat#A10491-01, Gibco) supplemented with 10% fetal bovine serum. The A549 and A375 cell lines were maintained in DMEM high glucose (Cat#08459-64, Nacalai Tesque) supplemented with 10% fetal bovine serum and in DMEM high glucose (Cat#08459-64, Nacalai Tesque) supplemented with sodium pyruvate and 10% fetal bovine serum, respectively. All cell lines were maintained at 37°C under 5% CO2.

### Genomic DNA preparation and sequencing

Genomic DNA was prepared from cells using QuickExtract solution (Nalgene) following the manufacturer’s instruction. PCR amplification was done using KOD FX Neo (Toyobo). PCR primers to amplify DNA were designed to target the oncogene mutated region (15^th^ exon for *BRAF* and 2^nd^ exon for *KRAS*). Direct sequencing of PCR products was carried out by FASMAC. The amplification primers were as follows: BRAF-Fw, 5’-ATCTCACCTCATCCTAACACATTTCAAGCCCC-3’; BRAF-Rv, 5’-GACTTTCTAGTAACTCAGCAGCATCTCAGGGCC-3’; KRAS -Fw, 5’-GCCTGCTGAAAATGACTGAA-3’; KRAS-Rv, 5’-AGAATGGTCCTGCACCAGTAA-3’.

### Soft-agar colony formation assay

Cells (2 × 10^4 per well) were mixed with 0.3% agarose, low gelling temperature (Cat#35640, SIGMA) in the full growth medium, plated on top of a solidified layer of 0.6% agarose in full growth medium ^41^ in a 6-well plate, and fed every 3 days with full growth medium. Photographs were taken by an OLYMPUS CKX53 inverted microscope with a DP20 digital camera (OLYMPUS). After 4 weeks, the colonies were stained with MTT (1 mg/ml in PBS solution) and imaged using an EPSON GT-X900 scanner.

### Western blot analysis

Cells were washed once with PBS and lysed directly in 1x SDS sample buffer (1 M Tris-HCl pH 6.8, 50% glycerol, 10% SDS, 0.2% bromophenol blue, and 10% 2-mercaptoethanol). After sonication and heat denaturation by boiling, the samples were separated by premade 5-20% gradient SDS-polyacrylamide gel electrophoresis (PAGE) (Nakalai or Atto) and transferred to Immobilon-FL Polyvinylidene Difluoride (PVDF) membranes (Millipore, Billerica, MA). After blocking with skim milk (Morinaga, Tokyo) or Odyssey blocking buffer (LI-COR), the membranes were incubated with primary antibodies diluted in skim milk, BSA, or Odyssey blocking buffer (LI-COR), followed by secondary antibodies diluted in Odyssey blocking buffer. Proteins were then detected by an Odyssey Infrared scanner (LI-COR) and analyzed by using the Odyssey software. For the analysis of ERK phosphorylation in the parental MCF-10A, KRAS G12V/+, and BRAF V600E/+ cell lines, cells were seeded with full growth medium for 1 day before the serum starvation; washed once with PBS and changed to starvation medium for 24 hrs; and finally treated with 10 ng/ml EGF for 10 min. In the case of ERK phosphorylation analysis in MCF-10A cells overexpressing KRAS G12V or BRAF V600E, the respective cell numbers were difficult to manipulate because of the different proliferation rates. Therefore, MCF-10A cells overexpressing KRAS G12V or BRAF V600E were seeded with the indicated medium, i.e., full growth medium or starvation medium, for 24 hrs and treated with 10 ng/ml EGF for 10 min.

### Oncogene-addiction experiments

For siRNA-mediated knockdown experiments to examine oncogene-addiction, we used siPOOLs (siTOOLs Biotech). The negative control siPOOL, BRAF targeted siPOOL and KRAS targeted siPOOL were purchased from siTOOLs Biotech. Reverse transfection of cancer cell lines and MCF-10A cell lines with siPOOLs was performed in Nunc edge 2.0 (96-Well Plates, Thermo Fisher) using Lipofectamine RNAiMax reagent (Invitrogen) according to the siPOOLs transfection protocol. Seeding density and siPOOL concentrations were modified depending on the cell lines and culture conditions used. After 3 to 4 days of siPOOL treatment, cells were washed once with PBS and stained with 0.1% crystal violet for 10 min, then washed three times with PBS and photographed with an EPSON GT-X900 scanner (1200 dpi). Relative cell number calculations were performed by ImageJ (Fiji) in comparison with the corresponding cell lines treated with a negative control siPOOL. For cancer cell lines, 1 nM siPOOL-reverse transfected cells were cultured in the RPMI 1640 medium (ATCC modification) (Cat#A10491-01, Gibco) supplemented with 10% fetal bovine serum for 4 days with the following cell densities, 6×10^3 cells/well for H358, 1×10^3 cells/well for A549, and 1-2×10^3 cells/well for A375. And knock-down experiment of the derivative cell lines of MCF-10A was performed the following conditions; in partial growth medium condition, reverse transfection in 6×10^3 cells with 1 nM siPOOLs for 3 days, and in starvation medium condition, reverse transfection in 3×10^3 cells with 0.5 nM siPOOLs for 4 days.

## Author contributions

R.E.I., C.O., and K.A. designed the research; R.E.I. performed the experiments and analyzed the data; and R.E.I., C.O., and K.A. wrote the paper.

## Acknowledgments

We thank Yusuke Mianari for the use of the bioruptor; Emi Ebine, and Kaori Onoda for their assistance; and all members of the K.A. laboratory for helpful discussions. This work was supported by the Functional Genomics Facility, NIBB Core Research Facilities. K.A. is supported by JST, CREST Grant No. JPMJCR1654; and by MEXT/JSPS KAKENHI Grant No. 16KT0069, 16H01425 “Resonance Bio”, 18H04754 “Resonance Bio”, 18H02444, and 19H05798.

## Conflict of Interest

The authors declare that they have no conflicts of interest with respect to the contents of this article.

## Notes

### Competing Interest Statement

The authors have declared no competing interest.

## References

1 Vogelstein B, Papadopoulos N, Velculescu VE, Zhou S, Diaz LA Jr, Kinzler KW. Cancer genome landscapes. Science 2013; 339: 1546–1558.

2 Hanahan D, Weinberg RA. Hallmarks of cancer: the next generation. Cell 2011; 144: 646–674.

3 Land H, Parada LF, Weinberg RA. Tumorigenic conversion of primary embryo fibroblasts requires at least two cooperating oncogenes. Nature 1983; 304: 596–602.

4 Ruley HE. Adenovirus early region 1A enables viral and cellular transforming genes to transform primary cells in culture. Nature 1983; 304: 602–606.

5 Watnick RS, Cheng Y-N, Rangarajan A, Ince TA, Weinberg RA. Ras modulates Myc activity to repress thrombospondin-1 expression and increase tumor angiogenesis. Cancer Cell 2003; 3: 219–231.

6 Weinstein IB. Cancer. Addiction to oncogenes--the Achilles heal of cancer. Science. 2002; 297: 63–64.

7 Weinstein IB, Joe A. Oncogene addiction. Cancer Res 2008; 68: 3077–80; discussion 3080.

8 Weinstein IB, Joe AK. Mechanisms of Disease: Oncogene addiction - A rationale for molecular targeting in cancer therapy. Nat Clin Pract Oncol 2006; 3: 448–457.

9 Sawyers CL. Opportunities and challenges in the development of kinase inhibitor therapy for cancer. Genes Dev 2003; 17: 2998–3010.

10 Sharma SV, Settleman J. Oncogene addiction: setting the stage for molecularly targeted cancer therapy. Genes Dev 2007; 21: 3214–3231.

11 Felsher DW. Oncogene addiction versus oncogene amnesia: perhaps more than just a bad habit? Cancer Res 2008; 68: 3081–6; discussion 3086.

12 Baylin SB, Ohm JE. Epigenetic gene silencing in cancer - a mechanism for early oncogenic pathway addiction? Nat Rev Cancer 2006; 6: 107–116.

13 Nishida E, Gotoh Y. The MAP kinase cascade is essential for diverse signal transduction pathways. Trends Biochem Sci 1993; 18: 128–131.

14 Cobb MH. MAP kinase pathways. Prog Biophys Mol Biol 1999; 71: 479–500.

15 Mukhopadhyay T, Tainsky M, Cavender AC, Roth JA. Specific inhibition of K-ras expression and tumorigenicity of lung cancer cells by antisense RNA. Cancer Res 1991; 51: 1744–1748.

16 Singh A, Greninger P, Rhodes D, Koopman L, Violette S, Bardeesy N et al. A Gene Expression Signature Associated with ‘K-Ras Addiction’ Reveals Regulators of EMT and Tumor Cell Survival. Cancer Cell 2009; 15: 489–500.

17 Hingorani SR, Jacobetz MA, Robertson GP, Herlyn M, Tuveson DA. Suppression of BRAF(V599E) in human melanoma abrogates transformation. Cancer Res 2003; 63: 5198–5202.

18 Sumimoto H, Miyagishi M, Miyoshi H, Yamagata S, Shimizu A, Taira K et al. Inhibition of growth and invasive ability of melanoma by inactivation of mutated BRAF with lentivirus-mediated RNA interference. Oncogene 2004; 23: 6031–6039.

19 Karasarides M, Chiloeches A, Hayward R, Niculescu-Duvaz D, Scanlon I, Friedlos F et al. B-RAF is a therapeutic target in melanoma. Oncogene 2004; 23: 6292–6298.

20 Sharma A, Trivedi NR, Zimmerman MA, Tuveson DA, Smith CD, Robertson GP. Mutant V599EB-Raf regulates growth and vascular development of malignant melanoma tumors. Cancer Res 2005; 65: 2412–2421.

21 Chin L, Tam A, Pomerantz J, Wong M, Holash J, Bardeesy N et al. Essential role for oncogenic Ras in tumour maintenance. Nature 1999; 400: 468–472.

22 Fisher GH, Wellen SL, Klimstra D, Lenczowski JM, Tichelaar JW, Lizak MJ et al. Induction and apoptotic regression of lung adenocarcinomas by regulation of a K-Ras transgene in the presence and absence of tumor suppressor genes. Genes Dev 2001; 15: 3249–3262.

23 Soule HD, Maloney TM, Wolman SR, Peterson WD Jr, Brenz R, McGrath CM et al. Isolation and characterization of a spontaneously immortalized human breast epithelial cell line, MCF-10. Cancer Res 1990; 50: 6075–6086.

24 Di Nicolantonio F, Arena S, Gallicchio M, Zecchin D, Martini M, Flonta SE et al. Replacement of normal with mutant alleles in the genome of normal human cells unveils mutation-specific drug responses. Proc Natl Acad Sci U S A 2008; 105: 20864–20869.

25 Konishi H, Karakas B, Abukhdeir AM, Lauring J, Gustin JP, Garay JP et al. Knock-in of mutant K-ras in nontumorigenic human epithelial cells as a new model for studying K-ras mediated transformation. Cancer Res 2007; 67: 8460–8467.

26 de Larco JE, Todaro GJ. Growth factors from murine sarcoma virus-transformed cells. Proc Natl Acad Sci U S A 1978; 75: 4001–4005.

27 Rothenberg SM, Engelman JA, L. S, Riese DJ 2nd, Haber DA, Settleman J. Modeling oncogene addiction using RNA interference. Proc Natl Acad Sci U S A 2008; 105: 12480–12484.

28 Hannus M, Beitzinger M, Engelmann JC, Weickert M-T, Spang R, Hannus S et al. siPools: highly complex but accurately defined siRNA pools eliminate off-target effects. Nucleic Acids Res 2014; 42: 8049–8061.

29 Lake D, Corrêa SAL, Müller J. Negative feedback regulation of the ERK1/2 MAPK pathway. Cell Mol Life Sci 2016; 73: 4397–4413.

30 Rambow F, Rogiers A, Marin-Bejar O, Aibar S, Femel J, Dewaele M et al. Toward Minimal Residual Disease-Directed Therapy in Melanoma. Cell 2018; 174: 843–855.e19.

31 Tsoi J, Robert L, Paraiso K, Galvan C, Sheu KM, Lay J et al. Multi-stage Differentiation Defines Melanoma Subtypes with Differential Vulnerability to Drug-Induced Iron-Dependent Oxidative Stress. Cancer Cell 2018; 33: 890–904.e5.

32 Khaliq M, Fallahi-Sichani M. Epigenetic Mechanisms of Escape from BRAF Oncogene Dependency. Cancers 2019; 11. doi: 10.3390/cancers11101480.

33 Banerji S, Cibulskis K, Rangel-Escareno C, Brown KK, Carter SL, Frederick AM et al. Sequence analysis of mutations and translocations across breast cancer subtypes. Nature 2012; 486: 405–409.

34 Cheng H, Liu P, Ohlson C, Xu E, Symonds L, Isabella A et al. PIK3CA(H1047R)-and Her2-initiated mammary tumors escape PI3K dependency by compensatory activation of MEK-ERK signaling. Oncogene 2016; 35: 2961–2970.

35 Liu Z, Wang L, Yang J, Bandyopadhyay A, Kaklamani V, Wang S et al. Estrogen receptor alpha inhibits senescence-like phenotype and facilitates transformation induced by oncogenic ras in human mammary epithelial cells. Oncotarget 2016; 7: 39097–39107.

36 Oikonomou E, Makrodouli E, Evagelidou M, Joyce T, Probert L, Pintzas A. BRAF(V600E) efficient transformation and induction of microsatellite instability versus KRAS(G12V) induction of senescence markers in human colon cancer cells. Neoplasia 2009; 11: 1116–1131.

37 Wajapeyee N, Serra RW, Zhu X, Mahalingam M, Green MR. Oncogenic BRAF induces senescence and apoptosis through pathways mediated by the secreted protein IGFBP7. Cell 2008; 132: 363–374.

38 Serrano M, Lin AW, McCurrach ME, Beach D, Lowe SW. Oncogenic ras provokes premature cell senescence associated with accumulation of p53 and p16INK4a. Cell 1997; 88: 593–602.

39 Tuveson DA, Shaw AT, Willis NA, Silver DP, Jackson EL, Chang S et al. Endogenous oncogenic K-rasG12D stimulates proliferation and widespread neoplastic and developmental defects. Cancer Cell 2004; 5: 375–387.

40 Jonkers J, Berns A. Oncogene addiction: sometimes a temporary slavery. Cancer Cell 2004; 6: 535–538.

41 Horibata S, Vo TV, Subramanian V, Thompson PR, Coonrod SA. Utilization of the Soft Agar Colony Formation Assay to Identify Inhibitors of Tumorigenicity in Breast Cancer Cells. J Vis Exp 2015; : e52727.

