## Supplemental Information for "Oncogenic mutation or overexpression of oncogenic KRAS or BRAF is not sufficient to confer oncogene addiction"

### Supplemental figure

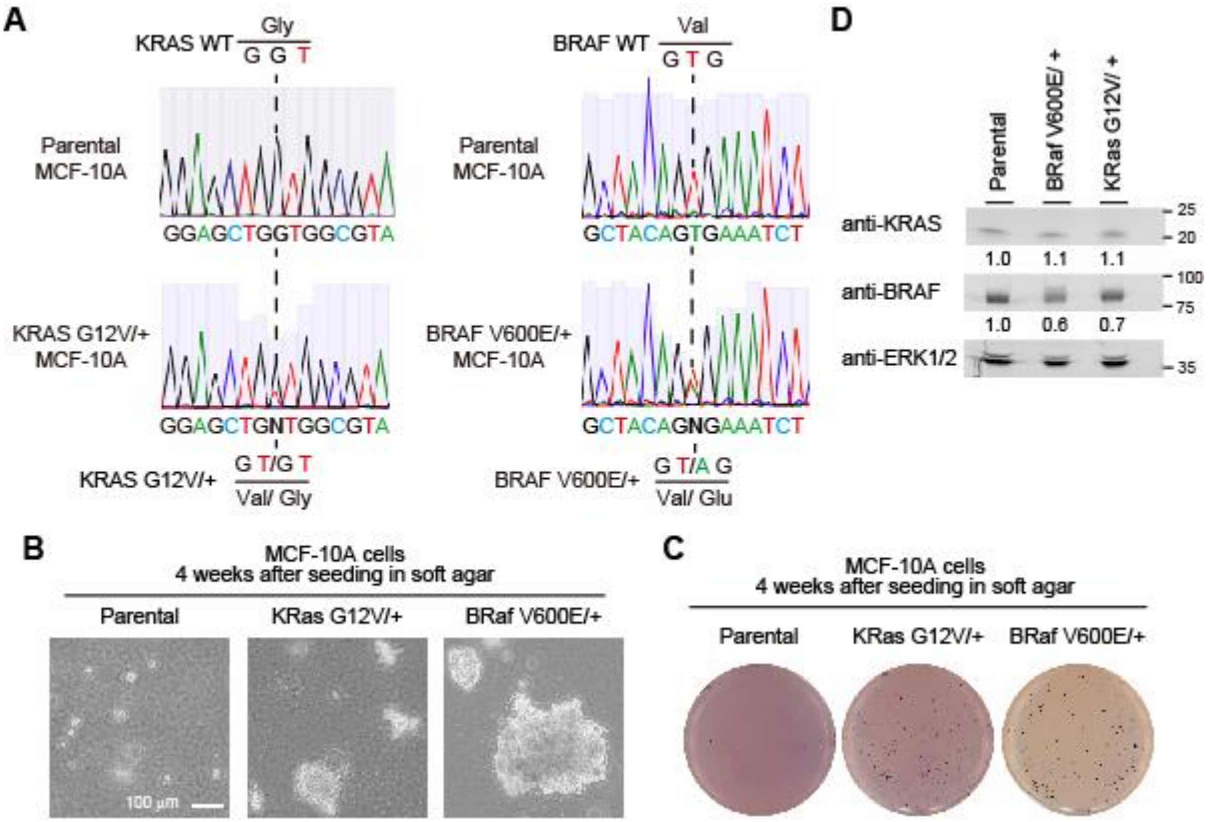


**Supplemental Figure 1. Characterization of MCF-10A cells harboring an oncogenic mutation in a single allele of KRAS or BRAF.**

(A) Genome sequence of *KRAS* gene (left) and *BRAF* gene (right) in the parental MCF-10A, KRAS G12V/+, and BRAF V600E/+ cell lines.

(B) Morphology of the indicated MCF-10A cells seeded in soft agar for 4 weeks.

(C) Representative images of MTT-stained colonies of the indicated MCF-10A cells seeded in soft agar for 4 weeks.

(D) Expression levels of KRAS and BRAF in the parental MCF-10A, KRAS G12V/+, and BRAF V600E/+ cell lines were analyzed by western blotting.

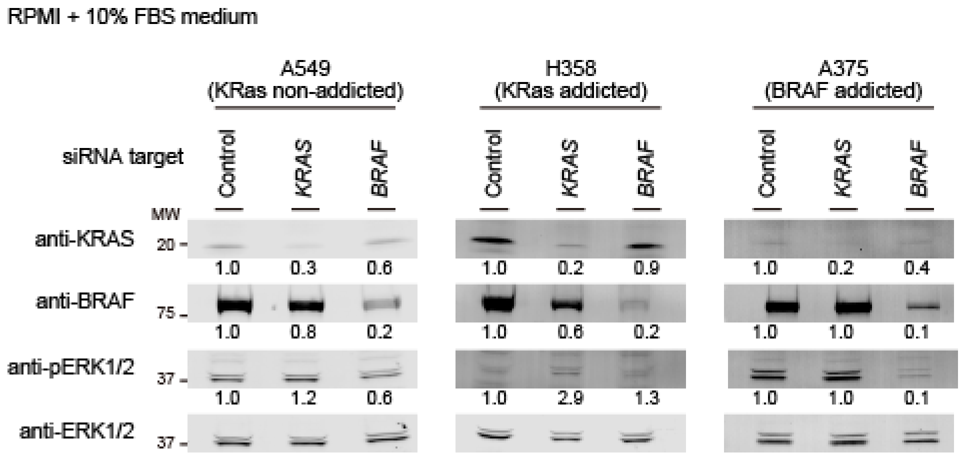


**Supplemental Figure 2. Knockdown efficiencies in cancer cell lines.**

Knockdown efficiencies of KRAS and BRAF using targeted siPOOLs in A549 cells (left), H358 cells (middle), and A375 cells (right) were analyzed by western blotting.

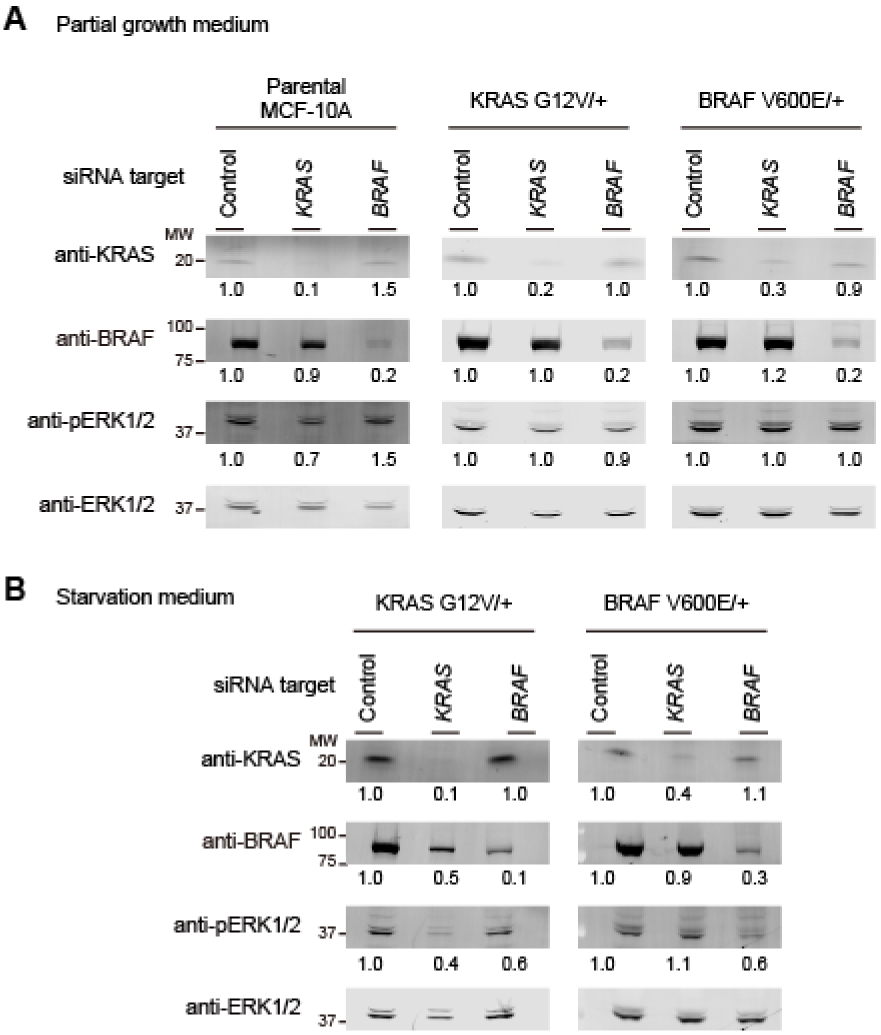


**Supplemental Figure 3. Knockdown efficiencies in MCF-10A cells harboring an oncogenic mutation in a single allele of *KRAS* or *BRAF*.**

Knockdown efficiencies of KRAS and BRAF using targeted siPOOLs in partial medium (A) and in starvation medium (B) were analyzed by western blotting.

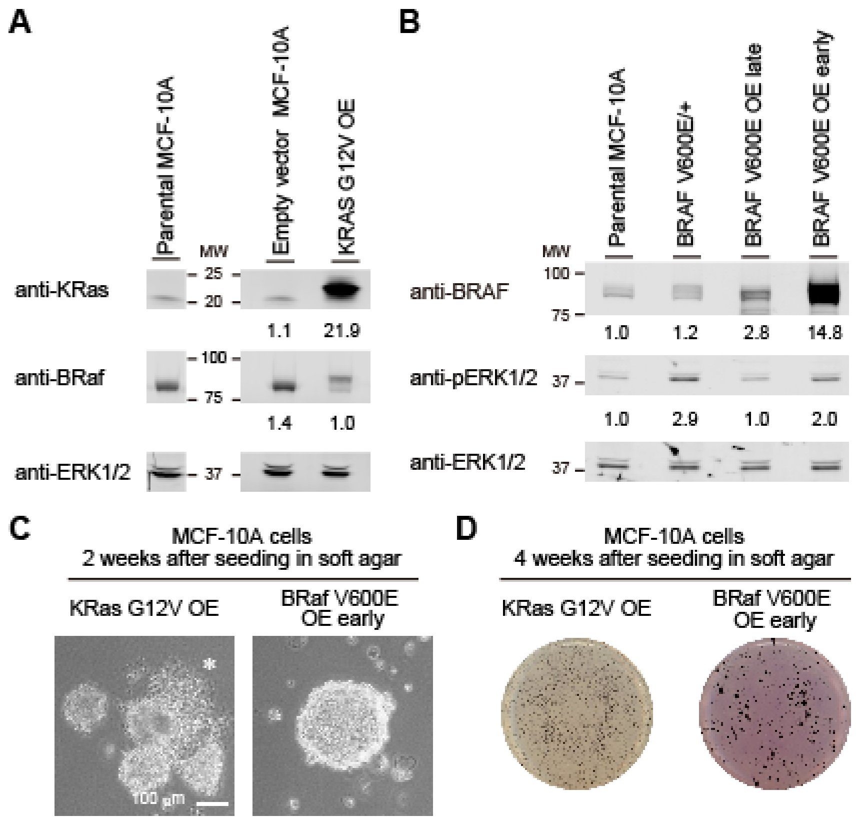


**Supplemental Figure 4. Characterization of MCF-10A cells overexpressing KRAS G12V or BRAF V600E.**

(A) KRAS expression levels of empty vector-introduced MCF-10A cells (control) and MCF-10A cells overexpressing KRAS G12V were analyzed by western blotting. The expression level was compared with that in parental MCF-10A cell lines.

(B) BRAF expression levels among the indicated cell lines were analyzed by western blotting. BRAF V600E OE early cells were established using pCSIIbsr-FLAG-BRaf V600E within 1 week.

(C) Morphology of the indicated MCF-10A cells seeded in soft agar for 2 weeks. A white asterisk indicates the ruptured spheroid in KRAS G12V OE.

(D) Representative images of MTT-stained colonies of the indicated MCF-10A cells seeded in soft agar for 4 weeks.

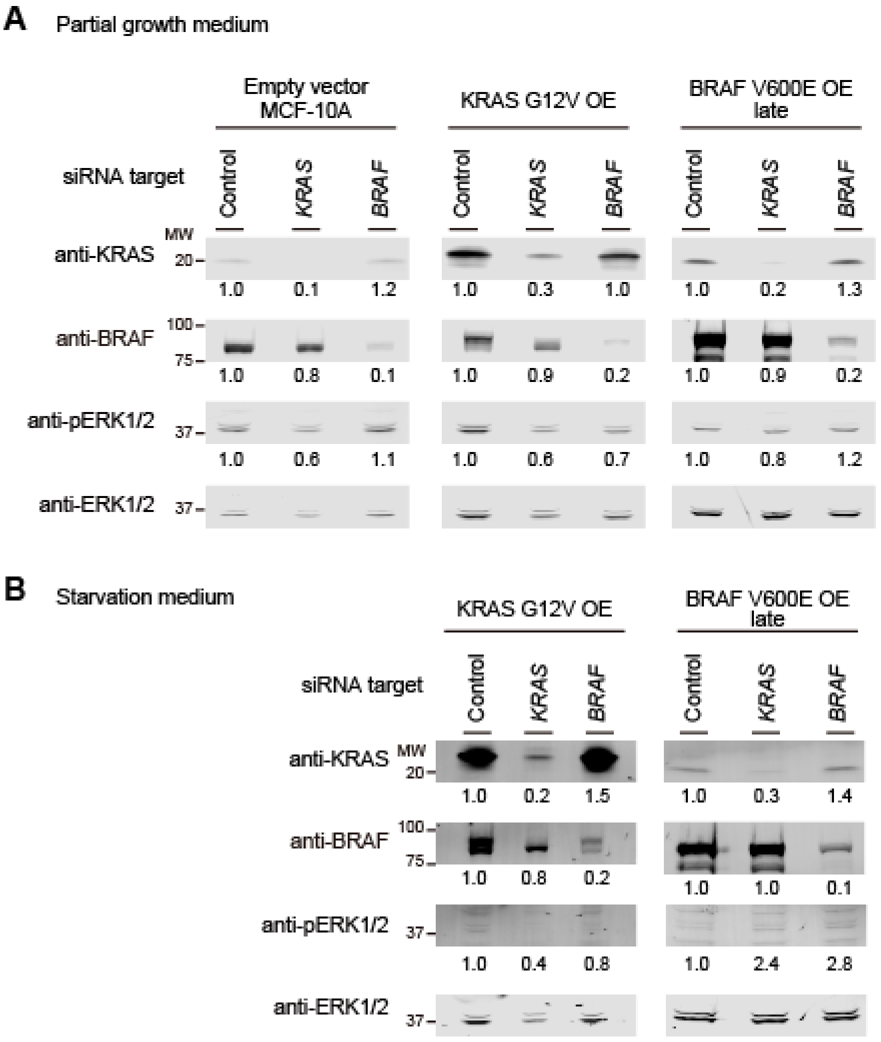


**Supplemental Figure 5. Knockdown efficiencies in MCF-10A cells overexpressing KRAS G12V or BRAF V600E.**

Knockdown efficiencies of KRAS and BRAF using targeted siPOOLs in partial medium (A) and in starvation medium (B) were analyzed by western blotting.
